# Cardiometabolic stress induces atrial myopathy in mice, characterized by structural remodelling, metabolic dysfunction, and incomplete transcriptional recovery

**DOI:** 10.64898/2026.10.06.754161

**Authors:** Jacques Couet, Diwaba Carmel Teou, Audrey Morin-Grandmont, Nicolas Le Bos, Thomas Le Bos, Emylie-Ann Labbé, Sara-Ève Thibodeau, Vincent Joseph, Marie Arsenault

## Abstract

**Background:** The mechanisms underlying atrial remodelling and recovery following the withdrawal of cardiometabolic stress remain incompletely understood. We investigated the functional, metabolic, and transcriptional consequences of metabolic-hypertensive stress (MHS), a murine model of cardiometabolic HFpEF, in the left atrium (LA) and assessed the reversibility of these alterations following myocardial recovery.

**Methods and Results:** Male and female mice underwent a two-hit HFpEF-inducing protocol consisting of angiotensin II infusion and a high-fat diet combined with physical inactivity (metabolic-hypertensive stress, MHS). Following 4 weeks of MHS, mice underwent a 4-week myocardial recovery (MR) phase involving stress withdrawal, dietary normalization, and voluntary exercise. We assessed LA function by speckle-tracking echocardiography, mitochondrial respiration by high-resolution respirometry, and transcriptomic remodelling by bulk RNA sequencing.

MHS markedly reduced LA reservoir, conduit, and contractile strain, consistent with atrial myopathy. Transcriptomic analyses revealed substantially greater remodelling in the LA than previously observed in the left ventricle under comparable experimental conditions. Hallmark pathway analysis showed activation of extracellular matrix remodelling and profibrotic signalling pathways, along with suppression of oxidative phosphorylation and fatty acid metabolism. Weighted gene co-expression network analysis identified a structural remodelling module enriched for extracellular matrix organization and angiogenesis pathways, as well as a bioenergetic module enriched for mitochondrial respiratory functions. Although inflammatory and structural remodelling pathways largely normalized following myocardial recovery, mitochondrial and metabolic programs exhibited incomplete restoration. Persistent transcriptional alterations were concentrated within a stress-responsive gene network enriched for adaptive cellular response pathways. In parallel, a discrete regulatory module displayed significant sex-dependent expression patterns.

**Conclusions:** Metabolic-hypertensive stress induces profound atrial functional, metabolic, and transcriptomic remodelling, consistent with the development of cardiometabolic atrial myopathy. While many stress-induced changes resolve following myocardial recovery, key bioenergetic and stress-responsive transcriptional programs persist, suggesting that cardiometabolic stress leaves a lasting molecular imprint within the atrium.

## Introduction

Heart failure with preserved ejection fraction (HFpEF) has emerged as the predominant form of heart failure worldwide and is increasingly recognized as a heterogeneous syndrome encompassing multiple phenotypes (1,2). Among these, the cardiometabolic phenotype, characterized by obesity, hypertension, insulin resistance, and systemic inflammation, accounts for a substantial proportion of affected patients. Recent evidence suggests that excess adiposity and adipose tissue-derived signalling pathways contribute directly to cardiac remodelling, promoting myocardial dysfunction through metabolic, inflammatory, and microvascular mechanisms (3–5).

The left atrium (LA) has gained increasing attention as a key determinant of cardiovascular performance in HFpEF. Beyond its role as a passive marker of diastolic dysfunction, the LA actively contributes to ventricular filling through its reservoir, conduit, and contractile functions. Impairment of these functions is associated with exercise intolerance, heart failure hospitalization, atrial fibrillation, and mortality. Accordingly, the concept of atrial myopathy has emerged to describe the structural, functional, and molecular alterations affecting the atrium independently of overt arrhythmias (6).

Experimental and clinical studies have demonstrated that cardiometabolic stress promotes atrial myopathy characterized by structural remodelling, fibrosis, inflammation, and impaired mechanical function. (6–11)

However, considerably less is known regarding the accompanying metabolic adaptations within the atrium (2,6). Emerging evidence suggests that mitochondrial dysfunction and altered substrate utilization may contribute to atrial disease progression (2,3). Yet, the relationship between atrial bioenergetics, transcriptomic remodelling, and atrial mechanical dysfunction remains poorly understood (6,10). Moreover, whether these alterations resolve after removal of the initiating stressor remains largely unexplored.

The reversibility of cardiac remodelling has major biological and clinical implications. Therapeutic interventions targeting obesity, hypertension, and metabolic dysfunction can improve systemic and ventricular phenotypes in HFpEF (12–14). Nevertheless, growing evidence suggests that prior cardiometabolic stress may leave persistent molecular signatures despite apparent functional recovery (15–17). Identifying which biological pathways normalize and which remain altered after stress withdrawal could provide important insights into the mechanisms underlying residual cardiovascular risk and disease recurrence (18–19).

In the present study, we used a two-hit murine model of cardiometabolic HFpEF induced by angiotensin II infusion and a high-fat diet combined with physical inactivity, referred to as metabolic-hypertensive stress (MHS) (20). After MHS, mice underwent a recovery phase characterized by withdrawal of the stressors, dietary normalization, and voluntary exercise (20,21). We hypothesized that cardiometabolic stress induces a distinct form of atrial myopathy associated with coordinated transcriptomic and mitochondrial remodelling, and that recovery would be incomplete despite normalization of several functional and molecular parameters. To test this hypothesis, we combined speckle-tracking echocardiography, high-resolution mitochondrial respirometry, and bulk RNA sequencing to characterize the left atrium’s functional, metabolic, and transcriptional landscape during disease induction and myocardial recovery.

We show that cardiometabolic stress triggers coordinated atrial mechanical dysfunction, mitochondrial remodelling, and transcriptomic reprogramming, with key bioenergetic and stress-responsive pathways remaining incompletely restored after myocardial recovery.

## Materials and Methods

### Animals

We used left atrial and left ventricular tissues from animals in a recently published investigation focused primarily on left ventricular remodelling and recovery following metabolic-hypertensive stress (MHS); we previously reported a detailed description of the experimental protocol (21). Each experimental group included eight male and eight female mice. See Figure M1 in the supplementary methods section for a schematic representation of the protocol.

### Metabolic-Hypertensive Stress (MHS)

As previously described (20,21), male and female mice received subcutaneous osmotic minipumps delivering angiotensin II (AngII; 1.5 mg/kg/day) for 28 days. Simultaneously, mice were fed either a high-fat diet (HFD; 60% kcal from fat) to induce MHS or a low-fat diet (LFD; 10% kcal from fat) as a control.

### Myocardial Recovery (MR)

After 28 days of MHS, osmotic minipumps were removed to terminate AngII infusion. Mice in the myocardial recovery (MR) group were switched from the HFD to the LFD. They were provided voluntary exercise through a flying saucer-type running wheel (Innowheel™, Innovive, Billerica, MA, USA) placed in their home cages. Animals remained under these recovery conditions for an additional 28 days, after which the protocol was completed.

We studied three experimental groups: control mice maintained on LFD (Ctrl), mice exposed to 28 days of metabolic-hypertensive stress (MHS), and mice studied after a 28-day myocardial recovery period following MHS (MR).

### Echocardiography

We performed echocardiography under isoflurane anesthesia (HR>450), as described previously (22,23). LA strain measurement. We measured LA strain using Vevo Strain speckle-tracking software (Vevo, Fuji Corp., Toronto, ON, Canada) according to a recently published method (24).

### Respirometry

#### Preparation of frozen cardiac tissue homogenates

We weighed frozen cardiac tissue and transferred it to a 1.5ml Eppendorf tube, kept on dry ice until needed for the experiment. The average tissue mass ranged from 0.6 to 1.0 mg for frozen mouse atrial tissue and from 0.2 to 0.6 mg for frozen ventricular tissue. We quickly thawed frozen tissue in 50 µL of mitochondrial respiration medium (MiR05) buffer and homogenized it using a hand-held tissue homogenizer. We loaded 50 µL of frozen tissue homogenate into the chambers of an Oroboros O2k with 2 mL of MiR05 buffer.

#### High-resolution respirometry of frozen cardiac tissue

We performed high-resolution respirometry under conditions similar to those reported in the literature (25,26) under normal oxygenated conditions (80–200 μM O_2_). We calculated mitochondrial respiratory capacity as O_2_ flux minus ROX, normalized to tissue wet weight, and expressed it as pmoles/s*mg tissue.

### Bulk mRNA-Sequencing

We created libraries from LA RNA using six individual samples per experimental group with the Illumina Stranded mRNA Library Prep Kit, following the manufacturer’s recommendations. We also analyzed libraries using the Agilent 2100 Bioanalyzer and DNA 1000 kit. We then quantified all samples using the KAPA Library Quantification Kit (Kapa Biosystems). We pooled the indexed libraries and sequenced them on the Illumina NovaSeq PE100 platform at Genome Quebec. We averaged 2.7 × 10^7 (1.4 to 3.7 × 10^7) paired-end reads, with Q30 of 39. We used Illumina BaseSpace RNA-Seq Alignment (STAR) for alignment and RNA-Seq Differential Expression (Dragen) to identify differentially expressed genes (DEGs) using the Mus musculus UCSC mm10 reference genome. We identified DEGs using an adjusted p-value ≤ 0.05 and a Log2 fold change (Log2FC) ≥ 0.5. The average library size was 289 base pairs. Gene set enrichment analysis was performed using the Panther GO Enrichment Analysis (geneontology.org) (27–29). We used a differentially expressed gene list (FDR < 0.05 and |Log2FC| ≥ 0.2 or ≥ 0.5) for this analysis. Significant GO terms were determined using Fisher’s Exact test with Benjamini–Hochberg correction with FDR.

### Hallmark gene set enrichment analysis (GSEA)

We performed GSEA to identify biological pathways associated with atrial and ventricular transcriptomic remodelling during metabolic-hypertensive stress (MHS) and myocardial recovery (MR). We conducted differential expression analyses using DESeq2, including sex as a covariate. We ranked genes by Wald statistics and analyzed them using Hallmark gene sets from MSigDB and the fgsea package (30–32). We quantified pathway activity using normalized enrichment scores (NES). To compare atrial and ventricular responses, we compared pathway-specific gene-level log2 fold changes between chambers. We calculated a recovery index to quantify the extent of pathway normalization after stress withdrawal. Additional methodological details are provided in the Supplementary Methods.

### Weighted Gene Co-expression Network Analysis (WGCNA)

We constructed gene co-expression networks using variance-stabilized RNA-sequencing data and the WGCNA package in R (33–35). We identified co-expression modules using a signed network approach and correlated them with experimental groups (CTRL, MHS, and MR). We characterized biologically relevant modules using Gene Ontology enrichment analyses. We integrated them with differential expression and Hallmark pathway analyses to identify transcriptional programs associated with cardiometabolic stress and myocardial recovery.

### Quantitative Real-Time Polymerase Chain Reaction

We used quantitative RT-PCR to quantify LV gene expression in at least six animals per group. Cyclophilin A (Ppia) was the control “housekeeping” gene. The primers used are listed in the Supplemental Methods section (Table M1).

### Statistical Analysis

All data are expressed as mean ± standard error of the mean (SEM). We removed outliers using the ROUT test (Q = 1%) in Prism. We conducted intergroup comparisons using a t-test in GraphPad Prism 11.1 (GraphPad Software Inc., La Jolla, CA, USA). We analyzed comparisons among more than two groups using one-way or two-way ANOVA (where appropriate) and a Holm-Sidak post hoc test. We considered P<0.05 statistically significant.

### Use of artificial intelligence tools

Artificial intelligence-assisted tools (Microsoft Copilot, GPT-5, Grammarly) supported the interpretation of transcriptomic datasets, including differential expression, gene ontology enrichment, Hallmark pathway analysis, and WGCNA results, as well as manuscript drafting and figure development. The authors independently verified all analytical workflows, statistical results, biological interpretations, and conclusions. The authors did not use AI tools to generate data or perform statistical computations.

## Results

### Metabolic-hypertensive stress (MHS) induces left atrial enlargement and alters ventricular structure and function

We previously investigated the animals included in the present study to better understand the determinants of LV recovery after MHS. These determinants were recovery duration (4 or 12 weeks), diet, and sedentarity (21). We weighed the left atria of these mice, then froze and stored them for further analysis. Here, we used these atrial tissues and associated echocardiographic data to examine the impact of MHS on atrial function and this chamber’s capacity to recover (MR).

Eight-week-old male and female mice received a four-week MHS regimen. This was associated with reduced body weight and tibial length in males. In females, an opposite trend was observed. After 4 weeks of recovery (16 weeks), when mice were fed a low-fat diet and allowed to exercise voluntarily, body weight and tibial length remained relatively unchanged (Figure 1A and B). As expected, the indexed heart (iHeart) and indexed LA (iLA) weights increased with MHS but returned to normal after recovery (Figure 1C and D).

**Figure 1.**
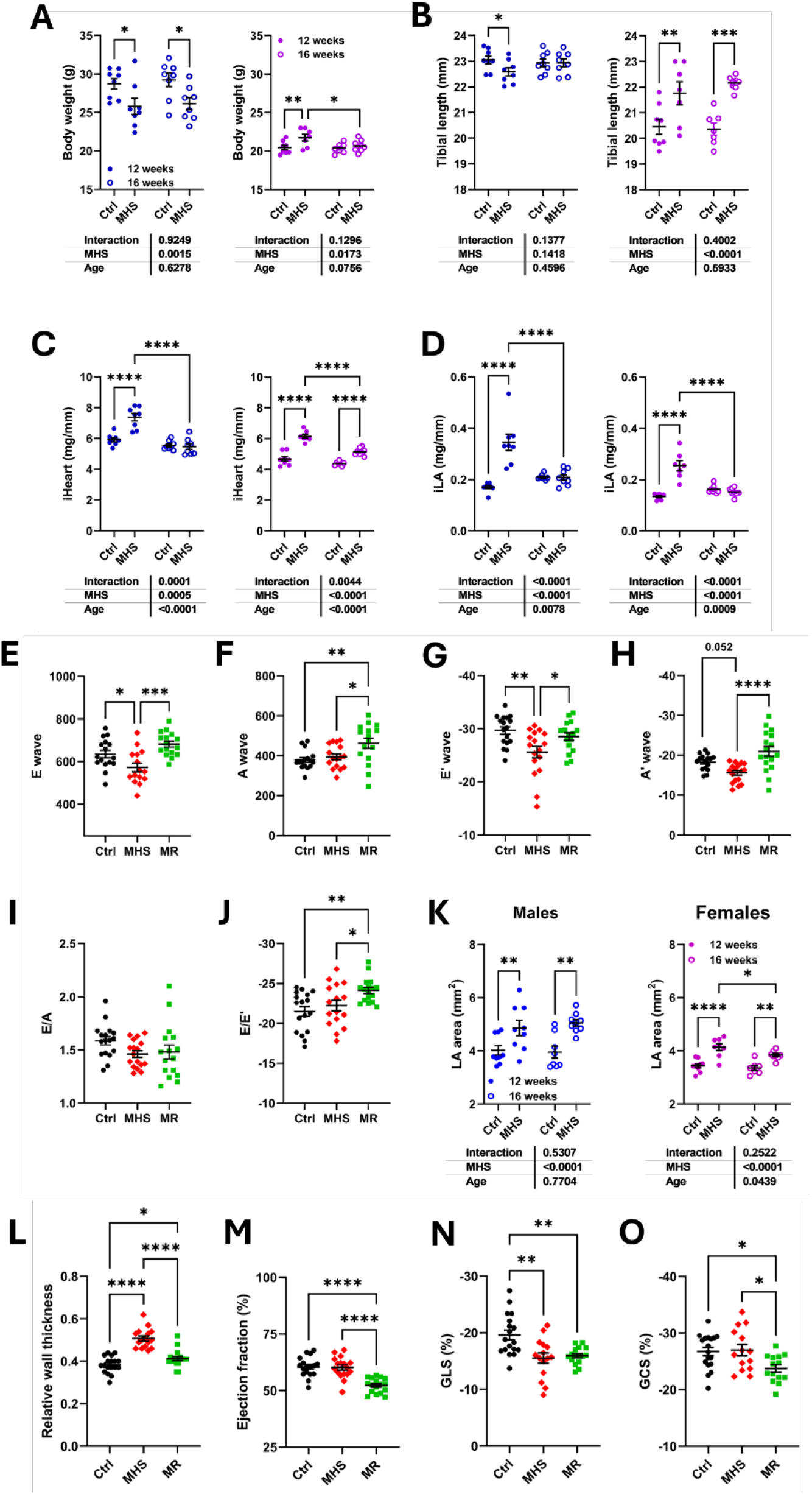
Morphometric and echocardiographic characterization of mice exposed to metabolic-hypertensive stress (MHS) and myocardial recovery (MR). (A) Body weight and (B) tibial length measured at 12 weeks (filled symbols) and 16 weeks of age (open symbols) in male (blue) and female (purple) mice. (C) Indexed heart weight (iHeart) and (D) indexed left atrial weight (iLA). Two-way ANOVA results for MHS, age, and interaction effects are shown below each panel. Echocardiographic assessment of left ventricular diastolic function: (E) mitral E-wave velocity, (F) mitral A-wave velocity, (G) tissue Doppler E′ velocity, (H) tissue Doppler A′ velocity, (I) E/A ratio, and (J) E/E′ ratio. (K) Left atrial (LA) area measured by echocardiography in male (blue) and female (purple) mice at 12 and 16 weeks of age. Indices of left ventricular geometry and systolic function: (L) relative wall thickness (RWT), (M) left ventricular ejection fraction (EF), (N) global longitudinal strain (GLS), and (O) global circumferential strain (GCS). Ctrl, control; MHS, metabolic-hypertensive stress; MR, myocardial recovery. Data are presented as mean ± SEM (n=14-16/gr). Panels E-O were analyzed using one-way ANOVA followed by Holm-Sidak post hoc testing. Panels A-D and K were analyzed using two-way ANOVA. *P < 0.05, **P < 0.01, ***P < 0.001, ****P < 0.0001.

The MHS regimen was also associated with alterations in LV diastolic function parameters, as illustrated in Figure 1E-J. Here, we studied male and female mice together. MHS reduced the pulsed-wave Doppler E wave and the tissue Doppler E’ wave. These two parameters were normalized 4 weeks after MHS cessation (MR). A and A’ waves were not modulated by MHS but were increased after MR. E/A and E/E’ ratios remained stable, except for the MR group (Figure 1J). LA area was estimated using echo (Figure 1K). MHS increased LA area, which remained larger than normal after MR.

We previously showed that the MHS regimen induced LV concentric remodelling (increased relative LV wall thickness, or RWT; Figure 1L) and that LV ejection fraction (EF; Figure 1M) was maintained as expected in an HFpEF model (20,21). Interestingly, LV global longitudinal strain was reduced in MHS animals (Figure 1N). After MR, RWT was mostly normalized, but EF was transiently reduced (Figure 1M). We previously observed that EF returned to normal after a longer recovery period of 12 weeks (21). Both GLS and GCS were below normal after MR, possible consistent with the transiently reduced EF (Figure 1N and O).

Although cardiac stressors were withdrawn during MR, indices of pulmonary congestion remained mildly elevated relative to controls (Supplementary Figure S1), suggesting incomplete physiological recovery.

### MHS mice have atrial myopathy as evidenced by reduced left atrial strain parameters

Reservoir strain corresponds to the maximal wall strain at the end of atrial diastole (Figure 2A). As shown in Figure 2B, the reservoir strain was reduced in MHS mice but returned to normal after 4 weeks of recovery (MR). Contractile strain corresponds to the LA deformation during atrial systole. In most mice with heart rates over 500 bpm, this becomes impossible to visualize, so we instead relied on the minimum LA strain rate as a marker of the start of atrial contraction to estimate contractile strain, as illustrated in Figure 2A. This point also corresponds to the P wave (atrial contraction) on the EKG. Contractile strain was reduced in MHS mice and returned to normal after 4 weeks of recovery (Figure 2B). Conduit strain, which can be interpreted as equivalent to the E wave, was also reduced in MHS mice, but in the MR group, it remained between controls and the MHS group (Figure 2B). By combining an index of LA mechanical function (LA reservoir strain) with an estimate of LV filling pressure (E/e′), the LA reservoir strain/E/e′ ratio has emerged as an integrated marker of atrial myopathy and diastolic dysfunction in humans and more recently, in mice (24,36). This ratio was increased in MHS animals and returned to normal in MR groups (Figure 5D). We illustrated sample LA strain tracings from one mouse per group (Figure 2C).

**Figure 2.**
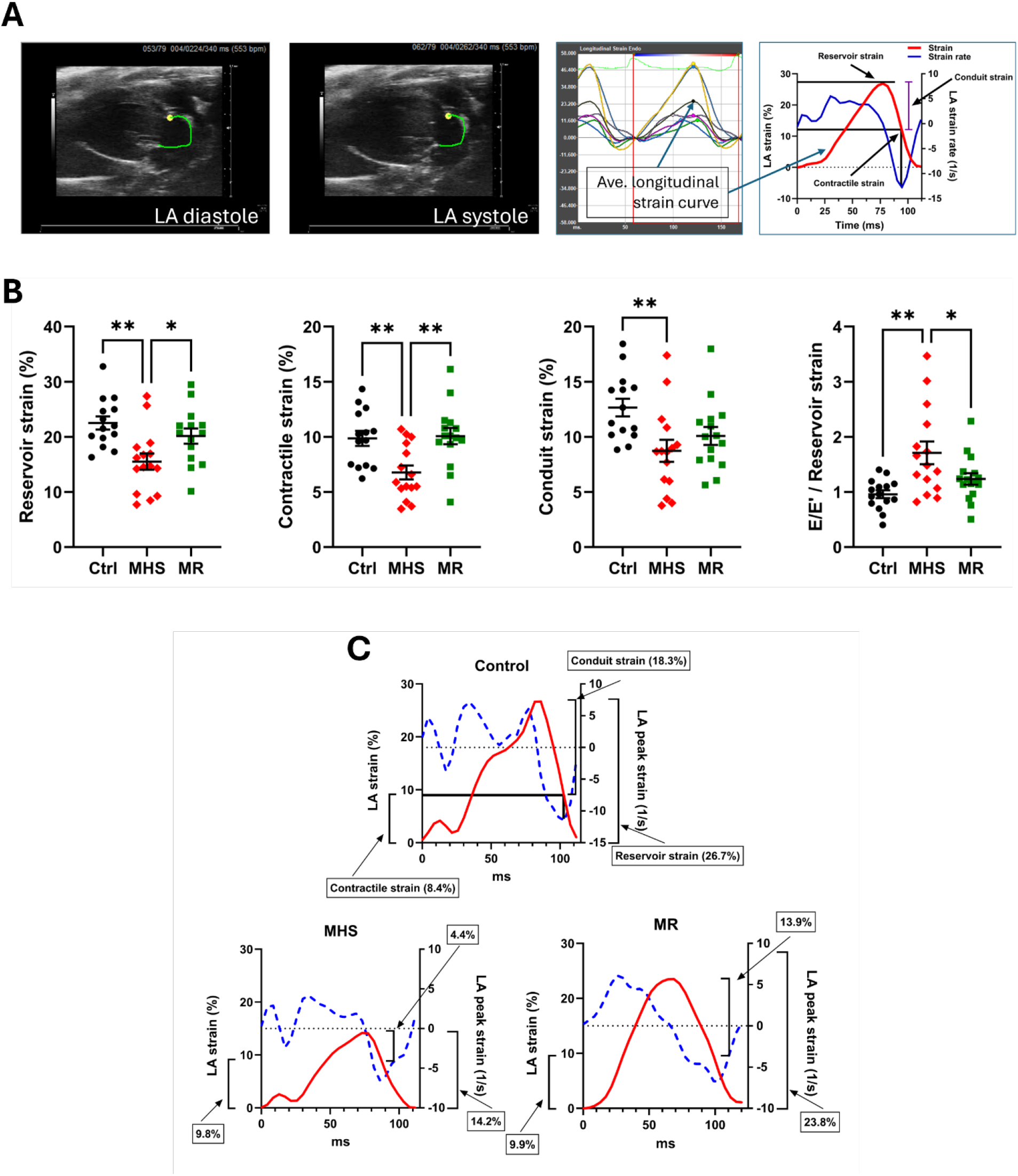
Metabolic-hypertensive stress impairs left atrial mechanical function and induces features of atrial myopathy. (A) Representative images illustrating left atrial (LA) strain analysis by speckle-tracking echocardiography. Endocardial border tracing of the LA in diastole and systole, average longitudinal strain assessment, and schematic representation of reservoir, conduit, and contractile strain measurements. We defined reservoir strain as peak LA strain during ventricular systole; conduit strain as the passive decline in strain during early ventricular filling; and contractile strain as the strain generated during atrial contraction, estimated from the minimum strain-rate signal because atrial and ventricular phases often fuse at high heart rates in mice. (B) Quantification of LA reservoir strain, contractile strain, conduit strain, and the E/E′ to reservoir strain ratio in control (Ctrl), metabolic-hypertensive stress (MHS), and myocardial recovery (MR) mice. MHS reduced all three components of LA strain and increased the E/E′/reservoir strain ratio. Recovery restored reservoir and contractile strain, whereas conduit strain remained intermediate between Ctrl and MHS groups. (C) Representative strain (red) and strain-rate (blue) tracings obtained from individual Ctrl, MHS, and MR mice, illustrating alterations in LA mechanical function during cardiometabolic stress and their partial normalization following recovery. Data are presented as mean ± SEM. One-way ANOVA followed by Holm-Sidak post hoc testing. *P < 0.05, **P < 0.01.

In a previous study (21), we had studied a longer recovery period of three months (12 weeks) after MHS (MR12). We measured LA strain parameters in these animals and compared them to age-matched mice (6-month-old). As shown in Figure S2, LA reservoir and contractile strains, as well as the E/E’/LA reservoir strain ratio, remained reduced in MR12 mice compared with controls, suggesting that LA function remained abnormal long after stress cessation.

### MHS mice have impaired atrial mitochondrial respiration

To determine whether MHS altered atrial bioenergetics, we measured mitochondrial respiration in LA homogenates using high-resolution respirometry (Figure 3). Basal oxygen consumption supported by malate, cytochrome c, and NAD⁺ was similar across experimental groups, indicating preserved basal respiratory activity (Figure 3A and 3B). In contrast, stimulation of complex I-linked respiration with acetyl-CoA revealed a significant reduction in oxygen flux in MHS animals compared with controls. Maximal complex I-supported respiration following NADH addition was similarly impaired during MHS, indicating a marked reduction in mitochondrial oxidative capacity. Following myocardial recovery, respiration partially improved but remained below control values. Adding succinate to stimulate convergent electron flow through complexes I and II increased oxygen consumption in all groups; however, MHS animals still showed a trend toward persistent impairment (Figure 3A and 3B). We then reproduced these measurements in the LV (Figure 3C). We did not observe this reduction in mitochondrial oxidative capacity after MHS, suggesting that LA mitochondrial respiration is more impaired after MHS than in the LV. MR mice showed a tendency for lower oxidative capacity that was not statistically significant. Together, these findings indicate that MHS reduces complex I-supported mitochondrial respiration in the LA but not the LV. Although MR tended to improve respiratory function in the LA, restoration remained incomplete, suggesting persistent bioenergetic impairment following withdrawal of cardiometabolic stress.

**Figure 3.**
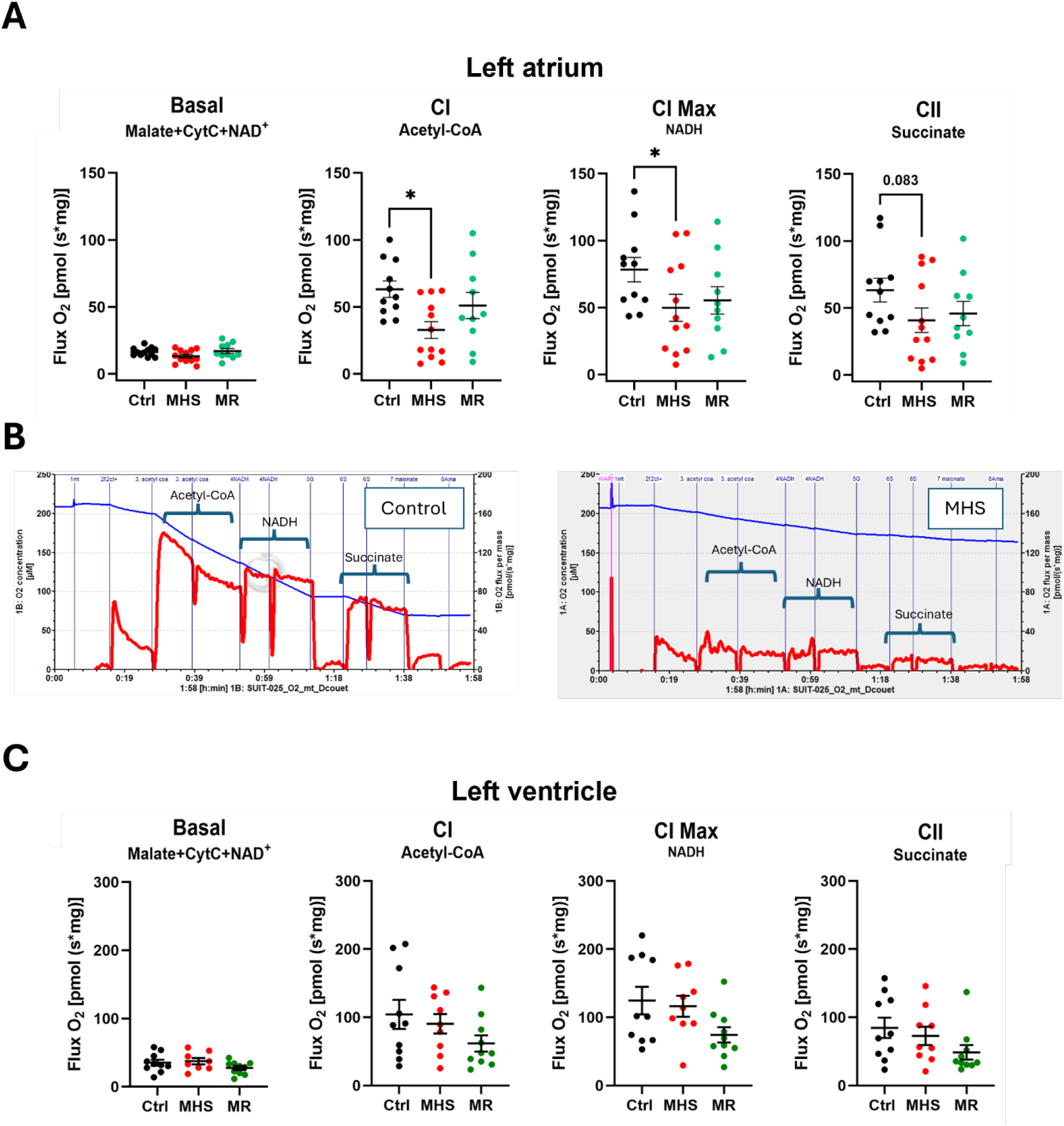
Metabolic-hypertensive stress impairs left atrial mitochondrial respiration. (A) High-resolution respirometry performed in left atrial homogenates from control (Ctrl), metabolic-hypertensive stress (MHS), and myocardial recovery (MR) mice. We measured basal oxygen flux in the presence of malate, cytochrome c, and NAD⁺. Complex I-supported respiration was assessed following sequential addition of acetyl-CoA (CI) and NADH (CI Max). We evaluated maximal convergent electron transport through complexes I and II after succinate addition (CII). (B) Representative oxygen flux traces obtained from a Ctrl mouse and an MHS mouse. Arrows indicate the sequential additions of acetyl-CoA, NADH, and succinate used to stimulate mitochondrial respiration. MHS markedly reduced complex I-supported respiration in the left atrium, as evidenced by lower oxygen flux following acetyl-CoA and NADH stimulation. Basal respiration was preserved, whereas respiration supported by convergent electron transport through complexes I and II showed a persistent trend toward impairment. Myocardial recovery partially restored respiratory capacity but did not fully normalize mitochondrial function. (C) High-resolution respirometry performed in left ventricular homogenates from control (Ctrl), metabolic-hypertensive stress (MHS), and myocardial recovery (MR) mice. Data are presented as mean ± SEM. LA: N=14-16/gr; LV: N=8-10/gr. One-way ANOVA followed by Holm-Sidak post hoc testing. *P < 0.05. Abbreviations: CI, complex I-supported respiration; CI Max, maximal complex I-supported respiration; CII, convergent complex I and II-supported respiration.

### Global transcriptomic remodelling is more pronounced in the left atrium than in the left ventricle after MHS

Taken together, the reductions in left atrial strain and mitochondrial respiratory capacity indicate that MHS induces both functional and metabolic components of atrial myopathy. We therefore performed bulk RNA sequencing to identify the transcriptional programs underlying these alterations and to determine how much they normalize following myocardial recovery.

Principal component analysis (PCA) revealed a clear separation between control, MHS, and MR samples in both cardiac chambers, indicating substantial transcriptomic remodelling induced by cardiometabolic stress and its subsequent withdrawal (Figure 4A and 4C). The separation was particularly pronounced in the LA, where the first PC explained 47.3% of total variance compared with 27.6% in the LV, suggesting a stronger chamber-specific transcriptional response to MHS. Consistent with these findings, differential expression analysis demonstrated markedly greater transcriptomic remodelling in the LA than in the LV. In pooled male and female analyses, MHS induced 2,496 differentially expressed genes (DEGs) in the LA compared with only 690 DEGs in the LV (Figure 4B, 4D and 4E). We observed similar differences when analyzing males and females separately.

**Figure 4.**
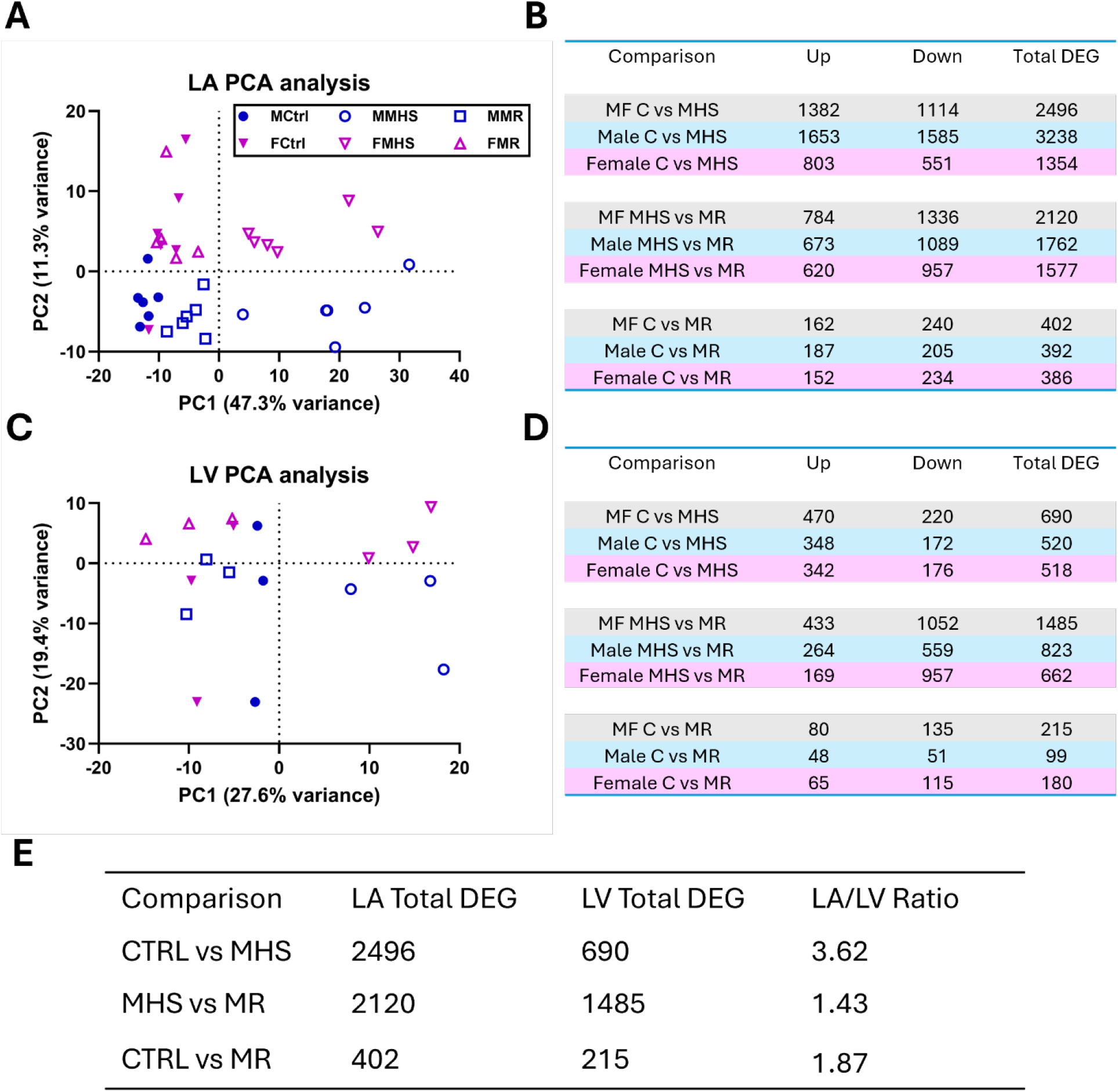
Cardiometabolic stress induces more extensive transcriptomic remodelling in the left atrium than in the left ventricle. (A, C) Principal component analysis (PCA) of left atrial (LA) and left ventricular (LV) transcriptomes from control (Ctrl), metabolic-hypertensive stress (MHS), and myocardial recovery (MR) mice. Male and female samples are displayed separately. In both chambers, MHS induced a marked shift in global gene expression that partially normalized following recovery. The separation between experimental groups was greater in the LA than in the LV, with PC1 accounting for 47.3% and 27.6% of total variance, respectively. (B, D) Numbers of differentially expressed genes (DEGs) identified in the LA and LV during MHS induction (Ctrl vs MHS), myocardial recovery (MHS vs MR), and residual remodelling after recovery (Ctrl vs MR). Results are shown for males and females separately and for both sexes combined. MHS induced substantially more transcriptomic remodelling in the LA than in the LV, whereas recovery was associated with extensive transcriptional reprogramming in both chambers. (E) Summary of chamber-specific transcriptomic responses. The LA exhibited 3.6-fold more DEGs than the LV during MHS (2496 vs. 690 genes), indicating greater sensitivity of the atrium to cardiometabolic stress. Although recovery markedly reduced the number of dysregulated genes, residual transcriptional alterations persisted in both chambers and remained more prominent in the LA than in the LV. DEG, differentially expressed gene; PC, principal component.

To validate chamber-specific remodelling patterns, we examined representative genes involved in extracellular matrix remodelling and mitochondrial metabolism using quantitative RT-PCR (Table S1). Structural remodelling markers, including *Ctgf, Ankrd1, Col1a1, Col3a1, Lox*, and *Mmp2*, were induced in both chambers but were generally more upregulated in the LA. In contrast, genes involved in mitochondrial respiration and energy metabolism, including *Cs*, *Ndufs1*, *Ndufb8*, *Sdha*, *Atp5fa1*, and *Tfam*, were more strongly suppressed in the LA, whereas ventricular changes were attenuated or absent. These data indicate that cardiometabolic stress elicits a conserved remodelling response in both chambers but a substantially more pronounced structural and metabolic reprogramming in the atrium. Modulation of additional genes in both chambers is shown in Table S2.

Given the substantially greater number of DEGs observed in the LA, we performed Hallmark pathway analysis to identify the biological processes underlying chamber-specific responses to MHS and MR (Tables 1 and S3). Although MHS elicited a broadly conserved remodelling program in both chambers, pathway-level analyses revealed markedly greater activation of extracellular matrix (ECM) remodelling, inflammatory signalling, and TGF-β pathways in the LA, together with a stronger suppression of oxidative phosphorylation (OXPHOS) and fatty acid metabolism. Hallmark Gene Set Enrichment Analysis (GSEA) also showed that after stress withdrawal, the LV robustly restored metabolic pathways, whereas the LA retained residual metabolic impairment.

### Myocardial recovery (MR) induces extensive transcriptional reprogramming

As mentioned above, following MHS withdrawal, MR was associated with substantial transcriptional reorganization in both chambers. Comparison of MR and MHS groups identified 2,120 DEGs in the LA and 1,485 DEGs in the LV (Figure 4B and D), indicating that recovery is an active biological process rather than a simple return toward the control state (Tables 1 and S5). Notably, the number of genes altered during MR approached the magnitude of changes induced during disease development, particularly in the LA, highlighting the dynamic nature of atrial remodelling in response to both MHS and MR.

### The MHS deregulates more LA genes in males than in females

Our RNA-Seq study included six males and six females (5 for MR). As shown in Figure 4B, when we split each group into two groups, MHS modulated more genes in males than in females. Males showed more than 500 additional LA genes modulated after MHS (3238 vs. 2496, whereas females had more than a thousand fewer (1354) than when both sexes were analyzed together. We performed a gene ontology (GO) analysis of male-specific upregulated and downregulated DEGs. Functional enrichment analysis showed that male-specific upregulated genes were predominantly associated with cytoskeletal remodelling, vesicular trafficking, intracellular signalling, and angiogenic processes (Table S4). In contrast, male-specific downregulated genes were enriched for pathways related to RNA metabolism, protein synthesis, mitochondrial organization, and fatty acid catabolism (Table S5).

### Weighted gene co-expression network analysis (WGCNA) identifies biologically distinct atrial remodelling programs

To identify coordinated transcriptional programs associated with LA remodelling and recovery, we performed WGCNA. Hierarchical clustering identified 14 co-expression modules, including the unassigned grey module (Supplementary Figures S3 and S4). Modules varied considerably in size and in their associations with atrial phenotype (Table S6). We prioritized five modules (turquoise, blue, black, red, and purple) for downstream analyses based on their significant correlations with left atrial weight and distinct eigengene dynamics across experimental groups (Figure 5A and Table 2).

**Figure 5.**
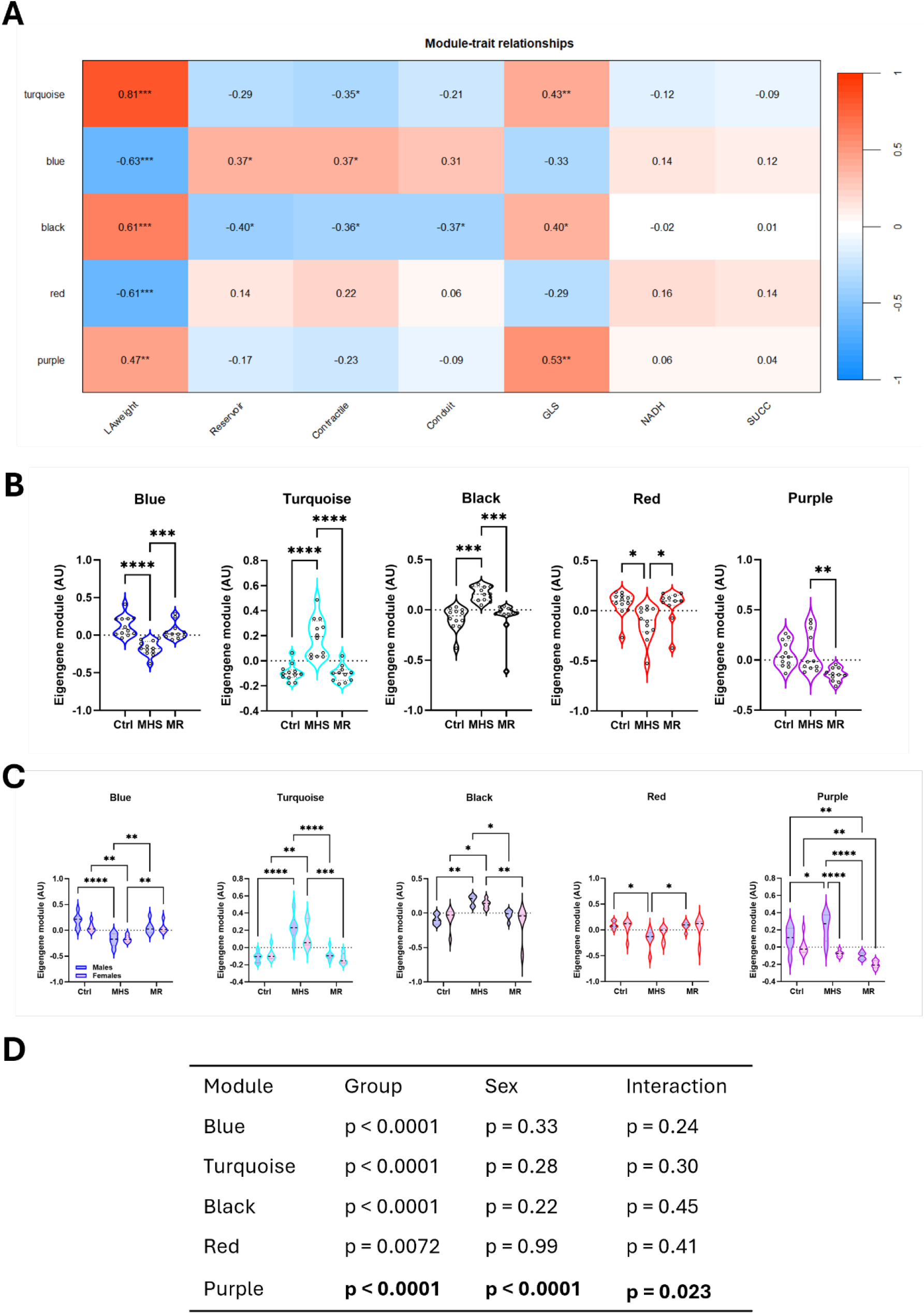
Weighted gene co-expression network analysis identifies distinct atrial remodelling, bioenergetic, and sex-dependent transcriptional programs. (A) Module-trait relationship heatmap showing correlations between eigengenes of the five principal WGCNA modules and phenotypic characteristics, including indexed left atrial weight (LA weight), left atrial strain parameters (reservoir, contractile, and conduit strain), global longitudinal strain (GLS), and mitochondrial respiratory capacity measured with NADH- and succinate-supported respiration. Red indicates positive correlations and blue indicates negative correlations. Correlation coefficients are displayed within each cell. P values: P < 0.05, P < 0.01, P < 0.001. (B) Module eigengene expression across experimental groups. The turquoise module (structural remodelling), black module (mechano signalling), and purple module (regulatory signalling) were induced during metabolic-hypertensive stress (MHS). In contrast, the blue module (mitochondrial bioenergetics) and red module (metabolic homeostasis) were suppressed. Most modules returned toward control levels following myocardial recovery (MR), whereas the purple module remained significantly altered. (C) Sex-stratified analysis of module eigengene expression. Responses of the blue, turquoise, black, and red modules were similar in males and females. In contrast, the purple module exhibited significant sex-dependent regulation across experimental groups. (D) Two-way ANOVA summary evaluating the effects of experimental group, sex, and the group-by-sex interaction on module eigengene expression. A significant interaction was observed only for the purple module, indicating sex-specific regulation of this transcriptional program. Data are presented as violin plots with individual animals shown as dots. Ctrl, control; MHS, metabolic-hypertensive stress; MR, myocardial recovery; GLS, global longitudinal strain; NADH, maximal complex I-supported respiration; SUCC, succinate-supported respiration. Module functional assignments were derived from Gene Ontology enrichment analyses: turquoise, structural remodelling; blue, mitochondrial bioenergetics; black, mechano signalling and cellular organization; red, metabolic homeostasis; purple, regulatory signalling and transcriptional control.

**Table 1.** Chamber-specific transcriptional responses to cardiometabolic stress and myocardial recovery. Hallmark gene set enrichment analysis (GSEA) revealed a conserved remodelling program in both cardiac chambers during metabolic-hypertensive stress (MHS). However, the left atrium (LA) showed greater activation of structural remodelling, profibrotic, and inflammatory pathways, along with stronger suppression of oxidative phosphorylation and fatty acid metabolism, than the left ventricle (LV). Following myocardial recovery (MR), inflammatory and remodelling pathways largely normalized, whereas metabolic pathways recovered incompletely in the LA, indicating persistent bioenergetic remodelling.

| Biological process | MHS response | Recovery |
| --- | --- | --- |
| OxPhos | More suppressed in LA | Restored in LV, incomplete in LA |
| FA metabolism | More suppressed in LA | Restored in LV, incomplete in LA |
| EMT/TGFβ | More activated in LA | Mostly normalized |
| Inflammation | More activated in LA | Resolved |

**Table 2.**
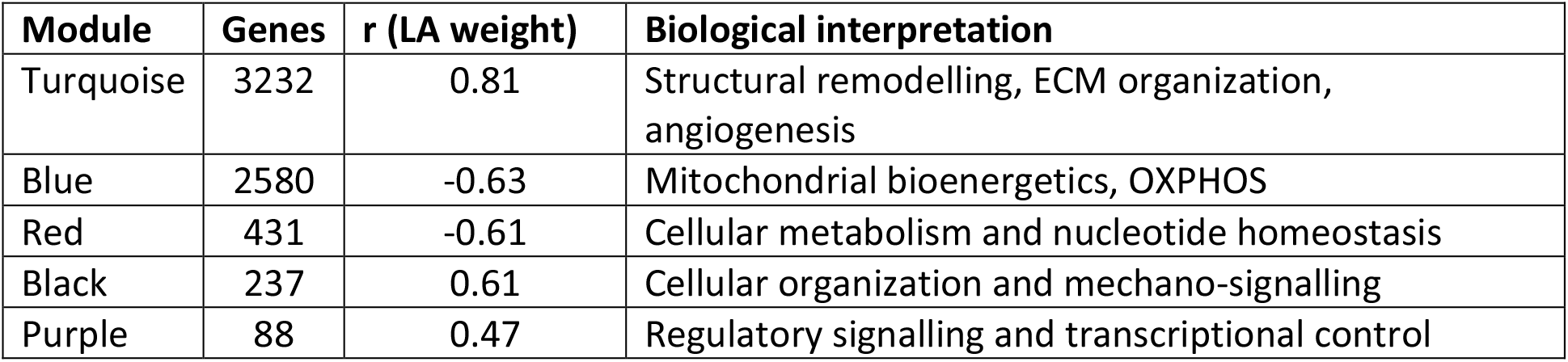
The five principal co-expression modules selected for downstream analyses are shown with their respective module size, correlation with indexed left atrial (LA) weight, and biological interpretation based on Gene Ontology enrichment analyses. The Turquoise and Black modules were positively associated with atrial remodelling, whereas the Blue and Red modules were inversely associated with atrial remodelling. The Purple module represented a regulatory transcriptional program exhibiting sex-dependent expression patterns.

| Module | Genes | r (LA weight) | Biological interpretation |
| --- | --- | --- | --- |
| Turquoise | 3232 | 0.81 | Structural remodelling, ECM organization, angiogenesis |
| Blue | 2580 | -0.63 | Mitochondrial bioenergetics, OXPHOS |
| Red | 431 | -0.61 | Cellular metabolism and nucleotide homeostasis |
| Black | 237 | 0.61 | Cellular organization and mechano-signalling |
| Purple | 88 | 0.47 | Regulatory signalling and transcriptional control |

The **turquoise module** (3232 genes) showed the strongest positive correlation with indexed LA weight (r = 0.81). It was enriched for extracellular matrix organization, angiogenesis, and structural remodelling pathways, identifying it as the principal atrial remodelling module. The **blue module** (2580 genes) exhibited a strong inverse association with atrial remodelling (r = −0.63) and was enriched for oxidative phosphorylation and mitochondrial respiratory pathways, consistent with a mitochondrial bioenergetic module. The **black module** (237 genes) was positively associated with atrial enlargement (r = 0.61) and was enriched for cellular organization and mechano-signalling pathways, suggesting a role in structural adaptation to MHS. The **red module** (431 genes) was inversely associated with atrial remodelling (r = −0.61) and was enriched for metabolic and nucleotide homeostasis pathways. Finally, the **purple module** (88 genes) exhibited a distinct expression pattern and was enriched for regulatory signalling and transcriptional control pathways, consistent with a sex-dependent regulatory module (Table S7).

### Distinct transcriptional modules exhibit differential recovery following stress withdrawal

Analysis of module eigengenes revealed distinct responses to cardiometabolic stress and myocardial recovery (Figure 5B). The turquoise and black modules, associated with LA structural adaptation and remodelling, were strongly induced during MHS and returned toward baseline following recovery, consistent with stress-dependent activation of structural remodelling and bioenergetic programs. Similarly, the blue (metabolism) and red (homeostasis) modules showed largely reversible responses after MHS withdrawal. In contrast, the purple module exhibited persistent suppression after MR, suggesting incomplete restoration of a discrete transcriptional program. When stratified by sex (Figure 5C), most module responses were remarkably similar in males and females. The notable exception was the purple module, which demonstrated significant sex-dependent regulation across experimental conditions, whereas stress-associated modules displayed comparable responses in both sexes.

### Persistent stress-responsive transcriptional programs remain after MR

Although MR markedly reduced atrial transcriptomic remodelling, comparison with Ctrl animals still identified 402 DEGs, indicating incomplete molecular recovery. To determine whether these residual alterations were concentrated in specific transcriptional networks, we mapped persistent genes onto WGCNA modules. Most persistent genes localized to the turquoise (n = 93) and blue (n = 133) modules, reflecting continued perturbations in structural remodelling and mitochondrial metabolic pathways (Table S8). However, the **GreenYellow** module exhibited the highest proportion of persistent genes (46% or 18/39 DEGs), suggesting enrichment for recovery-resistant transcriptional programs. Module eigengene analysis demonstrated that the GreenYellow module remained significantly suppressed following recovery, with no significant difference between MHS and MR groups (Figure 6A), indicating sustained dysregulation despite withdrawal of cardiometabolic stress. Representative genes within this network included the stress-responsive regulators *Dusp1, Dusp5, Klf2, Klf4, Nr4a2, Apold1, Ret,* and *Tnfaip6* (Figure 6B; Table S9). These genes are involved in adaptive cellular signalling, transcriptional regulation, endothelial stress responses, and ECM remodelling.

**Figure 6.**
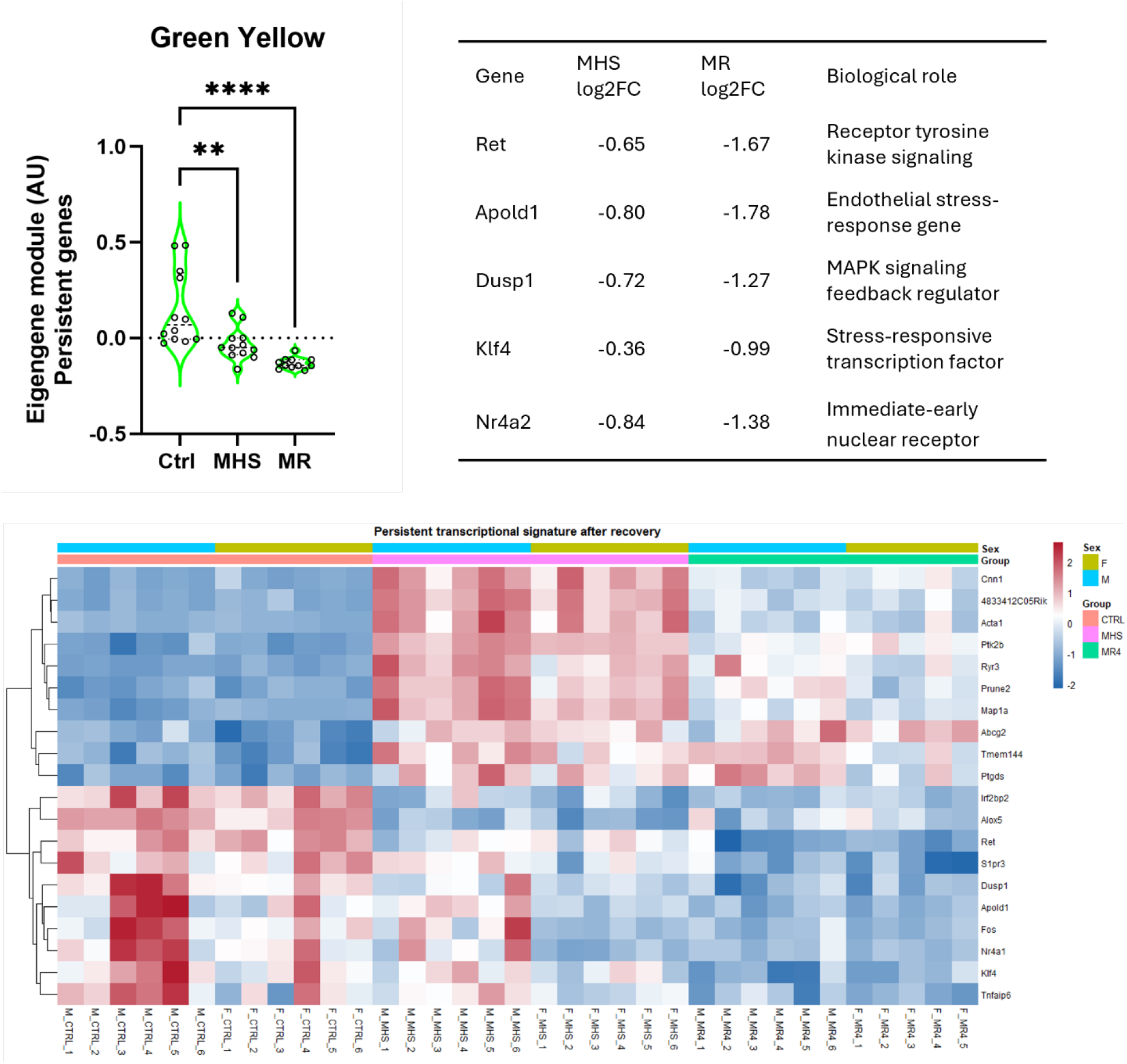
A stress-responsive transcriptional network exhibits persistent dysregulation following myocardial recovery. (A) Eigengene expression of the GreenYellow WGCNA module across control (Ctrl), metabolic-hypertensive stress (MHS), and myocardial recovery (MR) groups. The GreenYellow module contained the highest proportion of persistent genes following recovery (46.4%) and remained significantly suppressed in MR animals, with no significant difference between MHS and MR groups. (B) Representative persistent genes within the GreenYellow module. For each gene, log2 fold changes (log2FC) during MHS and after myocardial recovery relative to controls are shown together with the principal biological function. These genes are involved in stress-responsive signalling, transcriptional regulation, endothelial adaptation, and cellular remodelling. (C) Unsupervised hierarchical clustering of representative persistent genes from the GreenYellow module. Samples cluster primarily by experimental group rather than sex, demonstrating the persistence of a coordinated transcriptional signature after withdrawal of cardiometabolic stress. Representative genes include *Ret, Apold1, Dusp1, Fos, Nr4a1, Klf4, S1pr3*, and *Tnfaip6*, which remained dysregulated after recovery and collectively define a residual stress-response program within the left atrium. Data are presented as violin plots with individual animals shown as dots. log2FC, log2 fold change.

**Figure 7.**
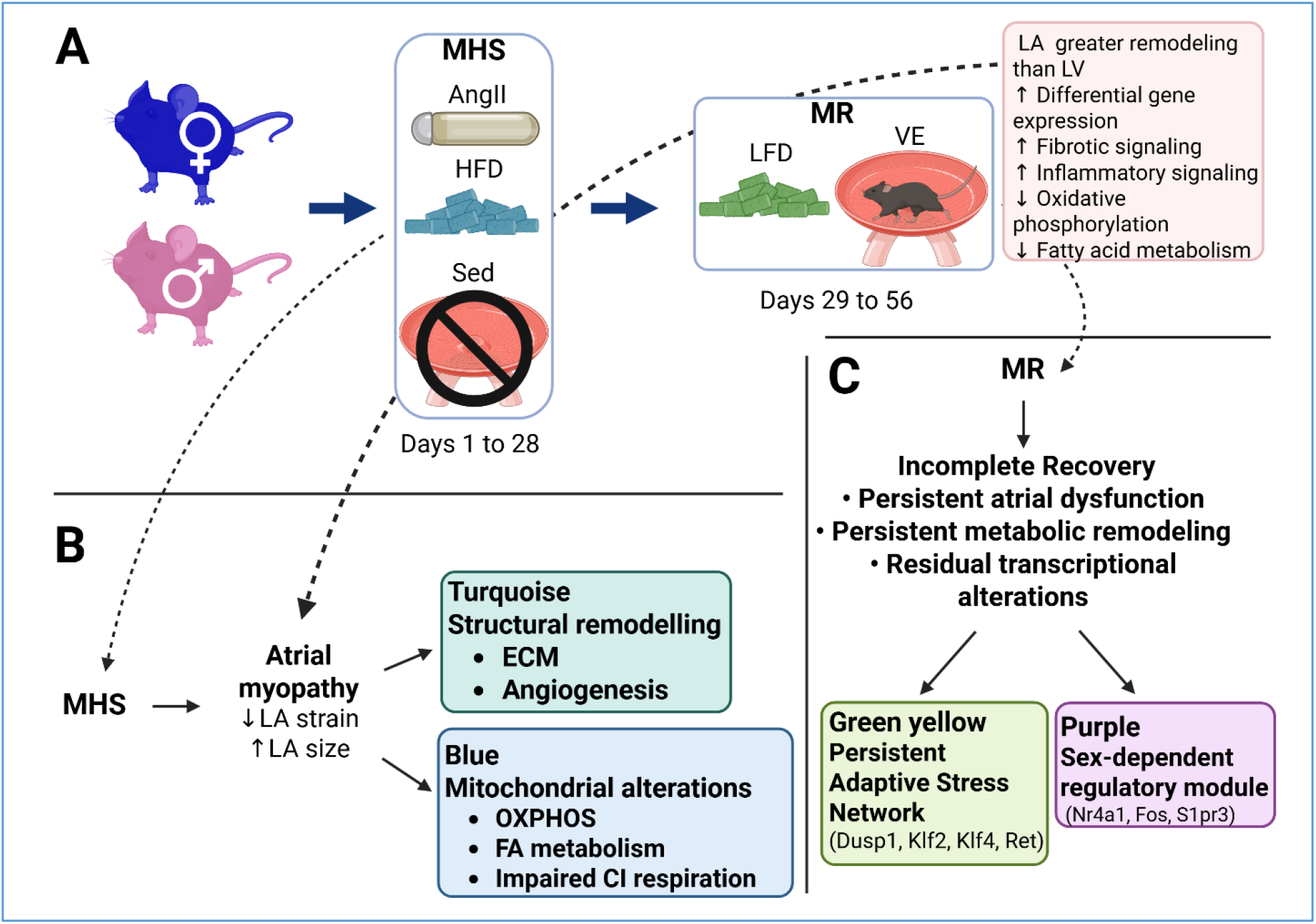
Integrated model of stress-induced atrial myopathy and transcriptional recovery following mechanical stress. (A) Experimental design. Mechanical stress (MHS) was induced by angiotensin II infusion, high-fat diet (HFD), and physical inactivity for 28 days. Recovery (MR) began with withdrawal of mechanical stress, replacement of the HFD with a low-fat diet (LFD), and voluntary exercise (VE) from day 29 to day 56. (B) Mechanical stress promotes an atrial myopathy phenotype characterized by impaired left atrial (LA) function, evidenced by reduced LA strain and increased LA size. Comparative transcriptomic analyses revealed a stronger atrial than ventricular response, with enhanced remodelling and fibrotic signalling together with suppression of oxidative phosphorylation (OXPHOS) and fatty-acid (FA) metabolism. WGCNA identified two major biological programs associated with atrial remodelling: a turquoise module enriched for extracellular matrix organization, angiogenesis, and structural remodelling pathways, and a blue module enriched for mitochondrial bioenergetic pathways, including OXPHOS, FA metabolism, and mitochondrial respiration. (C) Despite unloading, recovery remained incomplete. Persistent transcriptional alterations were enriched within the GreenYellow module, representing an adaptive stress-response network containing genes such as *Dusp1, Klf2, Klf4*, and *Ret*. In contrast, the purple module represented the principal sex-dependent regulatory network and was enriched for transcriptional and signalling pathways, including *Nr4a1, Fos*, and *S1pr3*. Together, these findings support a model in which mechanical stress induces structural remodelling and mitochondrial dysfunction in the atrium, with incomplete transcriptional recovery despite normalization of the initial stimulus. Created in BioRender. Couet, J. (2026) https://BioRender.com/fhr1tta.

Unsupervised clustering on persistent genes showed that samples remained segregated by prior MHS exposure rather than sex, supporting a persistent stress-response program within the atrium (Figure 6C). Together, these findings suggest that MHS leaves a durable molecular imprint characterized by incomplete normalization of adaptive transcriptional networks despite substantial physiological recovery.

## Discussion

### Atrial myopathy as a multidimensional disease process

In the present study, we used a young-mouse model of metabolic-hypertensive stress (MHS) induced by angiotensin II, high-fat diet feeding, and physical inactivity inspired by a similar HFpEF model in old female mice (37). Although this model differs from the more commonly used L-NAME/HFD paradigm (38), it recapitulates several key features of cardiometabolic HFpEF, including concentric ventricular remodelling, preserved ejection fraction, pulmonary congestion, impaired myocardial deformation, and atrial dysfunction (39). Using this model, we demonstrate that the left atrium undergoes profound functional, metabolic, and transcriptomic remodelling and exhibits incomplete recovery following withdrawal of cardiometabolic stress.

Recent work by Zhang MJ et al. (24) established left atrial strain as a robust marker of atrial myopathy in murine HFpEF models. Our findings extend this concept by showing that MHS induces global impairment of atrial function, characterized by reductions in reservoir, conduit, and contractile strain, an increased LA stiffness index, and atrial enlargement. Importantly, these alterations were accompanied by extensive metabolic and molecular remodelling, indicating that atrial myopathy reflects a complex biological process rather than an isolated mechanical abnormality (10,40). Collectively, the coexistence of impaired atrial mechanics, atrial enlargement, mitochondrial dysfunction, and transcriptomic remodelling supports the concept that MHS induces a bona fide atrial myopathy. Moreover, persistent LA enlargement and residual strain abnormalities after recovery suggest that this phenotype is only partially reversible.

Recent clinical and conceptual frameworks (11) have proposed that atrial myopathy may, in some contexts, develop as a primary disease process rather than simply reflecting ventricular dysfunction. While our findings do not establish this independence, the disproportionate functional, metabolic, and transcriptomic alterations observed in the left atrium relative to the left ventricle suggest that atrial remodelling exceeds ventricular remodelling and may reflect the atrium’s intrinsic susceptibility to cardiometabolic injury. Accordingly, atrial myopathy may be better viewed as a continuum of mechanical, metabolic, and molecular remodelling rather than a binary presence or absence of atrial dysfunction.

### The left atrium is particularly vulnerable to cardiometabolic stress

A major finding of the present study is that the left atrium appears substantially more sensitive to cardiometabolic stress than the left ventricle. The atrium exhibited more than three-fold more differentially expressed genes, stronger separation of experimental groups by principal component analysis, and more pronounced metabolic alterations. Hallmark analyses showed greater activation of extracellular matrix remodelling, EMT, TGF-β signalling, and inflammatory pathways, along with stronger suppression of oxidative phosphorylation and fatty-acid metabolism. These observations suggest that cardiometabolic stress elicits a broadly similar remodelling response in both chambers but a quantitatively amplified response in the atrium. This observation is consistent with recent clinical studies suggesting that atrial abnormalities may develop out of proportion to ventricular dysfunction in selected HFpEF phenotypes and are associated with adverse clinical outcomes, including hospitalization and atrial fibrillation (10, 40,41).

The basis for this chamber-specific vulnerability remains uncertain. The atrium is directly exposed to changes in ventricular filling pressures and operates under distinct mechanical constraints. However, the convergence of functional, metabolic, and transcriptomic findings consistently identifies the left atrium as a preferential target of cardiometabolic injury.

### Mitochondrial dysfunction as a central component of atrial remodelling

A particularly striking finding was the convergence of functional, transcriptomic, and network-level evidence implicating mitochondrial dysfunction as a central feature of atrial remodelling. High-resolution respirometry demonstrated marked impairment of complex I-supported respiration in the left atrium, whereas comparable impairments were not observed in the left ventricle. These findings were mirrored by suppression of oxidative phosphorylation and fatty-acid metabolism pathways and by the identification of the Blue WGCNA module enriched for mitochondrial respiratory functions.

Together, these observations suggest that impaired mitochondrial energetics is not simply a secondary consequence of atrial disease but a major biological component of the atrial response to cardiometabolic stress. Emerging evidence increasingly identifies mitochondrial dysfunction as a central mechanism linking cardiometabolic stress, systemic inflammation, energetic inefficiency, and cardiac dysfunction in HFpEF. Loss of metabolic flexibility, impaired oxidative phosphorylation, and altered substrate utilization have been proposed as fundamental determinants of disease progression (42–44). Although causality cannot be established, the parallel reduction in atrial strain, mitochondrial respiration, oxidative phosphorylation pathways, and mitochondrial gene expression supports a close association between bioenergetic failure and atrial dysfunction. Metabolic inflexibility is increasingly recognized as a hallmark of ventricular remodelling in HFpEF, our findings suggest that bioenergetic impairment may be even more pronounced in the left atrium, highlighting the possibility that atrial metabolic dysfunction contributes significantly to cardiometabolic atrial disease (43,44).

### Recovery is biologically selective rather than complete

Hallmark GSEA analyses suggest that structural and metabolic remodelling are tightly interconnected components of the atrial response to cardiometabolic stress. MHS activated extracellular matrix remodelling, EMT, TGF-β signalling, and inflammatory pathways while concurrently suppressing oxidative phosphorylation and fatty-acid metabolism. This pattern is consistent with a broad adaptive reprogramming in which the atrium shifts away from oxidative energy production while activating stress-response and tissue-remodelling pathways.

Importantly, recovery was not uniform across biological pathways. Inflammatory and structural signatures largely normalized following stress withdrawal, whereas suppression of oxidative phosphorylation and fatty-acid metabolism persisted. WGCNA further demonstrated that coherent transcriptional modules represented structural and bioenergetic programs. In contrast, the GreenYellow module identified a distinct recovery-resistant stress-response network containing genes involved in transcriptional regulation, endothelial signalling, and cellular adaptation. These findings suggest that myocardial recovery is biologically selective rather than complete and that prior cardiometabolic stress leaves a persistent molecular imprint within the atrium. This concept is consistent with the broader notion that reverse remodelling does not necessarily imply complete biological normalization and that residual molecular abnormalities may persist despite apparent physiological recovery (44).

### Clinical implications

A key strength of the present model is its ability to study biological recovery after near-complete removal of cardiometabolic stressors. Such conditions are rarely achieved in clinical practice, where obesity, hypertension, aging, and metabolic dysfunction often persist despite treatment. Our findings suggest that normalization of the initiating stimulus may not fully restore atrial biology and that apparent functional recovery may mask persistent biological vulnerability.

If the left atrium is disproportionately affected by cardiometabolic stress, longitudinal assessment of atrial structure and function may provide a sensitive indicator of disease progression and recovery. Persistent abnormalities in LA strain may identify patients with ongoing biological remodelling despite clinical improvement. Recent clinical studies suggest that atrial imaging markers may detect disease progression before overt chamber enlargement occurs and may provide prognostic information regarding future cardiovascular outcomes (10,41).

Our findings also suggest that therapies targeting mitochondrial function and metabolic flexibility could complement conventional approaches aimed at correcting hemodynamic and cardiometabolic stressors. However, the persistence of bioenergetic abnormalities after stress withdrawal suggests that the therapeutic window for complete recovery may be limited once remodelling is established. Several therapeutic strategies currently under investigation in HFpEF, including SGLT2 inhibitors, metabolic modulators, exercise-based interventions, and mitochondria-targeted approaches, have been proposed to improve mitochondrial function and energetic efficiency (44,45).

Although the present study did not investigate atrial fibrillation, several features observed in MHS mice, including atrial enlargement, impaired atrial mechanics, extracellular matrix remodelling, metabolic dysfunction, and persistent stress-responsive signalling, are commonly associated with an arrhythmogenic substrate (10,40). Whether atrial myopathy represents a consequence of HFpEF, a contributor to disease progression, or both remains uncertain. Nevertheless, our findings support the emerging concept that persistent atrial remodelling may have clinical relevance beyond atrial fibrillation alone and could contribute to the long-term cardiovascular burden associated with cardiometabolic disease (10,40).

### Study limitations

Several limitations should be considered when interpreting the present findings. First, this study was performed in a preclinical mouse model of cardiometabolic HFpEF. Although the metabolic-hypertensive stress model reproduces several important features of the human syndrome, including preserved ejection fraction, ventricular remodelling, pulmonary congestion, atrial dysfunction and decreased exercise capacity (20), it cannot fully capture the complexity, duration, and heterogeneity of HFpEF in patients. In addition, animals were studied at a young age. In contrast, HFpEF predominantly affects older individuals and is often influenced by aging-related biological processes not represented in the present model.

Second, the present model combines angiotensin II infusion, high-fat feeding, and physical inactivity but does not produce marked obesity because the catabolic effects of angiotensin II partially offset weight gain. Consequently, the relative contribution of obesity-related mechanisms to atrial remodelling may differ from that observed in patients with severe cardiometabolic HFpEF. Furthermore, we did not specifically investigate the contribution of sex hormones or menopause-related hormonal changes, which may be particularly relevant given the high prevalence of HFpEF among postmenopausal women. Although we included both sexes, the study was not designed to dissect the specific roles of gonadal hormones.

Third, chamber comparisons should be interpreted with some caution. Left ventricular RNA-sequencing data were derived from a related cohort studied previously (20), although several representative ventricular genes were independently validated in the current animals by quantitative PCR and demonstrated similar chamber-specific patterns (21).

Fourth, technical constraints imposed by the mouse’s small left atrium prevented simultaneous assessment of all endpoints in individual animals. Consequently, we could not measure fibrosis, transcriptomic profiling, and mitochondrial respiration in the same specimens, precluding direct correlations among histological, bioenergetic, and transcriptomic parameters. In addition, we assessed mitochondrial respiration using frozen tissue homogenates. While this approach allowed robust evaluation of oxidative capacity, it prevented direct assessment of substrate-specific metabolism using fatty acids, pyruvate, or glucose and therefore limited mechanistic characterization of atrial metabolic flexibility.

Fifth, network-based analyses should be considered exploratory and hypothesis-generating. Although WGCNA identified biologically coherent modules that were strongly associated with atrial phenotype and supported by independent pathway analyses, module composition and trait associations may be influenced by sample size and trait selection. The relatively modest number of animals available for RNA sequencing may therefore limit detection of weaker co-expression relationships and reduce the stability of some modules. Nevertheless, the convergence between differential expression analyses, Hallmark pathway enrichment, mitochondrial phenotyping, and WGCNA supports the overall robustness of the principal biological conclusions.

Sixth, we did not assess atrial electrophysiology or susceptibility to atrial arrhythmias. Consequently, the potential relationship between the observed molecular alterations and arrhythmogenic risk remains speculative.

Finally, the observational nature of the study does not allow us to determine whether mitochondrial dysfunction is a primary driver of atrial remodelling or a secondary consequence of cardiometabolic stress. Future studies using targeted metabolic interventions will be required to establish causality.

## Conclusion

In summary, metabolic-hypertensive stress induces a bona fide atrial myopathy characterized by coordinated mechanical, metabolic, and transcriptomic remodelling. The left atrium appears disproportionately vulnerable to cardiometabolic injury compared with the left ventricle, with mitochondrial dysfunction emerging as a prominent feature of this response. Although many functional and structural alterations improve following stress withdrawal, persistent metabolic and stress-responsive transcriptional abnormalities suggest that cardiometabolic stress leaves a lasting molecular imprint within the atrium.

## Grants

The Canadian Institutes of Health Research (PJT-1665850; J. Couet) and the Fondation de l’Institut universitaire de cardiologie et de pneumologie de Québec supported this work.

## Acknowledgements

The authors thank Mrs Élisabeth Walsh-Wilkinson for performing surgeries and echocardiographic exams and Mr. François Marcouillier for technical support with respirometry experiments.

