## Supplemental data for "Cardiometabolic stress induces atrial myopathy in mice, characterized by structural remodelling, metabolic dysfunction, and incomplete transcriptional recovery"

Jacques Couet, Diwaba Carmel Teou, Audrey Morin-Grandmont, Nicolas Le Bos, Thomas Le Bos, Emylie-Ann Labbé, Sara-Ève Thibodeau, Vincent Joseph, Marie Arsenault.

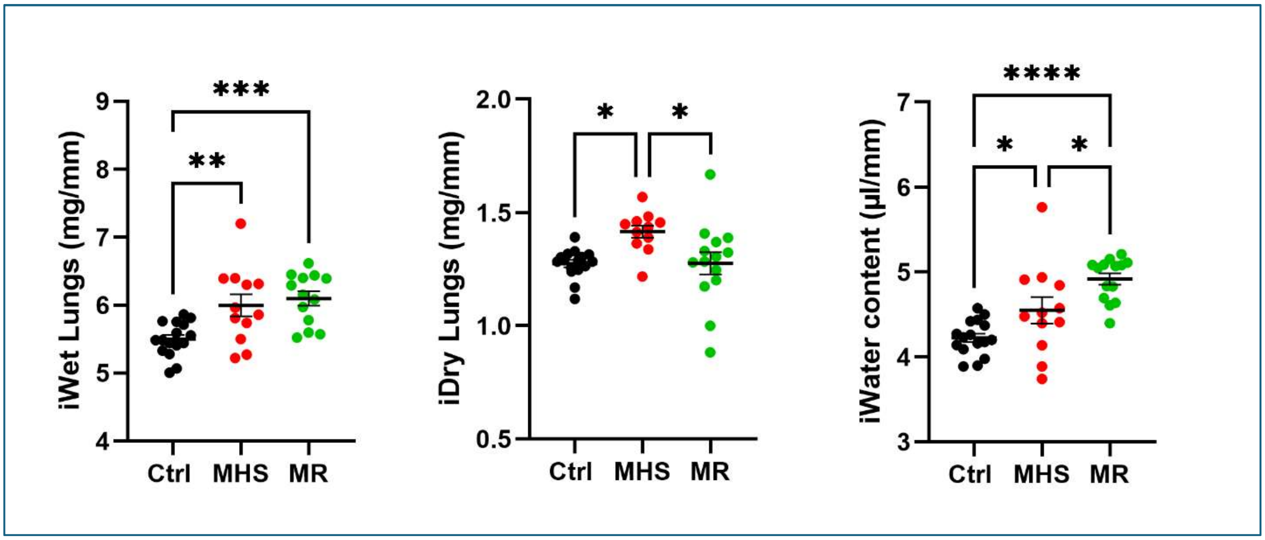


**Figure S1.** Indexed (for tibial length) lungs' wet and dry weights and water content (Wet - Dry). Results are shown as the mean ± SEM. One-way ANOVA followed by Holm-Sidak post-test. *: p<0.05, **: p<0.01, ***: p<0.001 and ****: p<0.0001 between indicated groups.

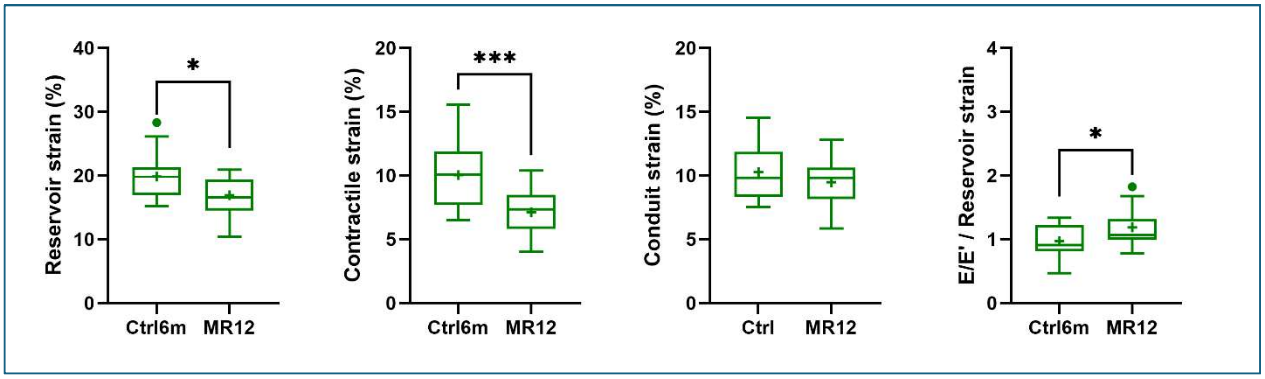


**Figure S2.** Reservoir, contractile, and conduit LA strains, as well as the E/E'/Reservoir strain ratio, in 6-month-old control animals (Ctrl) and mice after 12 weeks of MHS withdrawal (MR12). Results are shown as the mean ± SEM. One-way ANOVA followed by Holm-Sidak post-test. *: p<0.05, **: p<0.01, ***: p<0.001 and ****: p<0.0001 between indicated groups (8 males and 8 females/gr).

**Table S1.** Validation of chamber-specific (LA and LV) remodelling and metabolic signatures induced by MHS using quantitative RT-PCR. Results are indicated in terms of fold change over corresponding controls. (n=10-12 mice/gr). P values were calculated using Welch's t-test. NS: not significant.

| **Remodelling-associated genes** | | | | |
|---|---|---|---|---|
| **Gene** | **LA MHS/CTRL** | **p value** | **LV MHS/CTRL** | **p value** |
| *Ctgf/Ccn5* | 15.1 | <0.001 | 2.62 | <0.001 |
| *Ankrd1* | 7.22 | <0.001 | 2.77 | <0.001 |
| *Postn* | 4.98 | 0.0016 | 4.26 | <0.001 |
| *Col1a1* | 3.58 | <0.001 | 2.23 | 0.011 |
| *Col3a1* | 3.28 | <0.001 | 1.80 | 0.003 |
| *Lox* | 3.53 | 0.004 | 2.36 | <0.001 |
| *Mmp2* | 3.56 | <0.001 | 1.87 | 0.002 |

| **Mitochondrial and metabolic genes** | | | | |
|---|---|---|---|---|
| **Gene** | **LA MHS/CTRL** | **p value** | **LV MHS/CTRL** | **p value** |
| *Cs* | 0.63 | 0.001 | 0.93 | NS |
| *Ndufs1* | 0.62 | <0.001 | 0.92 | NS |
| *Ndufb8* | 0.70 | <0.001 | 0.93 | NS |
| *Sdha* | 0.61 | <0.001 | 0.82 | 0.006 |
| *Atp5fa1* | 0.63 | <0.001 | 0.85 | 0.047 |
| *Tfam* | 0.74 | 0.006 | 0.94 | NS |
| *Pink1* | 0.67 | <0.001 | 0.75 | 0.001 |

**Table S2**—a list of additional genes tested in the LA and LV after MHS. Results are indicated in terms of fold change over corresponding controls. (n=10-12 mice/gr). P values were calculated using Welch's t-test.

| Gene | MHS, LA, FC | p value | MHS, LV, FC | p value |
|---|---|---|---|---|
| *Nppa* | 1.87 | <0.0001 | 3.00 | 0.0002 |
| *Nppb* | 4.32 | 0.0036 | 2.49 | 0.0009 |
| *Myh6* | 0.95 | 0.60 | 0.88 | 0.010 |
| *Myh7* | 0.94 | 0.53 | 4.62 | 0.0068 |
| *Actc1* | 1.77 | 0.0005 | 0.87 | 0.33 |
| *Acta2* | 2.57 | 0.0020 | 1.47 | 0.0032 |
| *Tgfb3* | 2.29 | 0.0002 | 1.37 | 0.072 |
| *Ctsk* | 0.85 | 0.0005 | 1.74 | 0.0014 |
| *Idh3* | 0.67 | 0.0029 | 0.96 | 0.66 |
| *Uqcrc2* | 0.67 | 0.0021 | 0.94 | 0.32 |
| *Opa1* | 0.67 | 0.0028 | 0.87 | 0.12 |

**Table S3.** Hallmark gene set enrichment analysis (GSEA) of chamber-specific (LA vs. LV) transcriptomic responses to mechanical stress and recovery.

| Pathway | MHS Response (LA vs. LV) | Recovery (MR4) (LA vs. LV) | Biological Interpretation |
|---|---|---|---|
| **Oxidative Phosphorylation** | Stronger suppression in LA (NES: -3.31 vs -2.48; $P$=1.4×10$^{-30}$) | Restored in LV (NES: +3.06) but remained suppressed in LA (NES: -1.96; $P$=1.3×10$^{-28}$) | Persistent mitochondrial dysfunction and incomplete metabolic recovery in the LA |
| **Fatty Acid Metabolism** | Stronger suppression in LA (NES: -2.27 vs -1.46; $P$=6.7×10$^{-10}$) | Recovered in LV (NES: +1.97) but remained reduced in LA (NES: -0.96; $P$=2.8×10$^{-5}$) | Persistent impairment of lipid utilization in the LA |
| **TGF-beta** | Greater activation in LA (NES: 2.08 vs 1.50; $P$=6.0×10$^{-6}$) | Suppressed in both chambers, with stronger reversal in LV ($P$=0.045) | Enhanced fibrotic/remodeling response in LA during stress |
| **EMT** | Greater activation in LA (NES: 3.20 vs 2.58; $P$=3.6×10$^{-14}$) | Reversed in both chambers (NS at MR4) | Stronger structural remodelling in LA during MHS |
| **Inflammatory Response** | Greater activation in LA (NES: 2.00 vs 1.43; $P$=1.5×10$^{-3}$) | Largely resolved in both chambers (NS at MR4) | Stress-induced inflammation resolves after unloading |

**Table S4.** Biological processes enriched among male-specific upregulated genes following MHS. We performed Gene Ontology (GO) Biological Process overrepresentation analysis using the PANTHER classification system (version 19.0). We selected representative, non-redundant GO terms to summarize the major biological themes enriched among genes significantly upregulated in males following MHS. We assessed statistical significance using Fisher's exact test with false discovery rate (FDR) correction. Fold enrichment represents the observed-to-expected ratio for each GO term.

| **Biological theme** | **Representative GO term** | **Fold enrichment** | **FDR** |
|---|---|---|---|
| Cytoskeletal remodeling | Actin cytoskeleton organization (GO:0030036) | 3.61 | $1.97 \times 10^{-18}$ |
| Vesicular and endosomal trafficking | Endosomal transport (GO:0016197) | 3.68 | $1.57 \times 10^{-9}$ |
| Intracellular signaling | Intracellular signal transduction (GO:0035556) | 2.18 | $1.20 \times 10^{-8}$ |
| Cell migration and adhesion | Cell migration (GO:0016477) | 2.40 | $2.44 \times 10^{-6}$ |
| Phosphoinositide signaling | Phosphatidylinositol metabolic process (GO:0046488) | 3.13 | $1.44 \times 10^{-5}$ |
| Angiogenesis and vascular remodelling | Angiogenesis (GO:0001525) | 4.70 | $1.91 \times 10^{-5}$ |
| Cellular morphogenesis | Neuron development (GO:0048666) | 2.19 | $1.07 \times 10^{-4}$ |

**Table S5.** Biological processes enriched among male-specific downregulated genes following MHS. We performed Gene Ontology (GO) Biological Process overrepresentation analysis using the PANTHER classification system (version 19.0). We selected representative, non-redundant GO terms to summarize the major biological themes enriched among genes significantly downregulated in males following MHS. We assessed statistical significance using Fisher's exact test with false discovery rate (FDR) correction. Fold enrichment represents the observed-to-expected ratio for each GO term.

| **Biological theme** | **Representative GO term** | **Fold enrichment** | **FDR** |
|---|---|---|---|
| RNA and nucleic acid metabolism | Nucleic acid metabolic process (GO:0090304) | 2.01 | $1.95 \times 10^{-11}$ |
| Gene expression and biosynthesis | Gene expression (GO:0010467) | 1.65 | $1.92 \times 10^{-6}$ |
| Protein synthesis and translation | Translation (GO:0006412) | 2.70 | $1.86 \times 10^{-5}$ |
| RNA processing and splicing | RNA splicing (GO:0008380) | 2.69 | $3.80 \times 10^{-5}$ |
| Mitochondrial organization | Mitochondrion organization (GO:0007005) | 2.84 | $2.13 \times 10^{-5}$ |
| Fatty acid catabolism and β-oxidation | Fatty acid beta-oxidation (GO:0006635) | 7.57 | $4.79 \times 10^{-8}$ |
| DNA maintenance and repair | DNA damage response (GO:0006974) | 2.29 | $1.01 \times 10^{-4}$ |
| Immune response (underrepresented) | Immune response (GO:0006955) | 0.15 | $1.14 \times 10^{-11}$ |

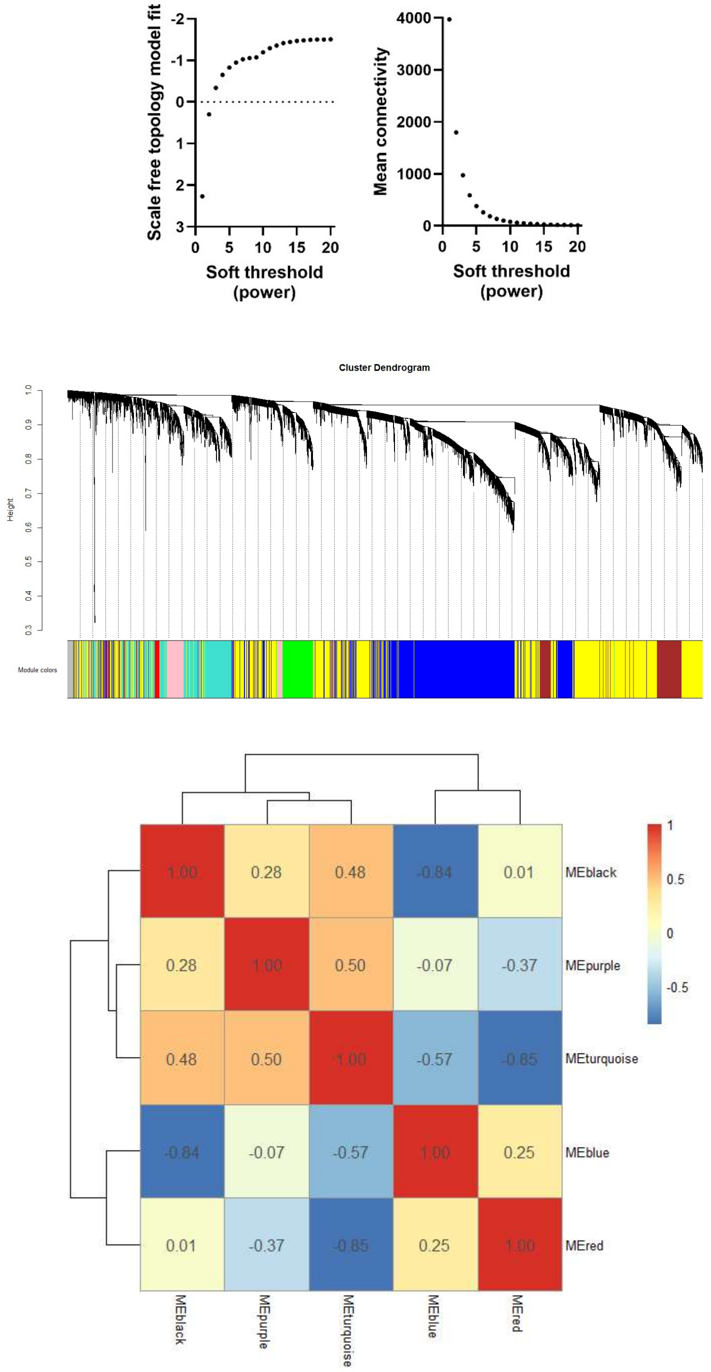


**Figure S3**. Construction and characterization of the weighted gene co-expression network. (up) Scale-free topology analysis was used to determine the optimal soft-thresholding power (β = 9) for network construction. This power produced a near scale-free topology while preserving adequate network connectivity. (middle) Hierarchical clustering dendrogram of 12,810 genes based on topological overlap dissimilarity. Colored bars indicate module assignments identified using dynamic tree cutting. We detected fourteen modules, including the unassigned grey module. (bottom) Hierarchical clustering of module eigengenes from the five modules selected for downstream analyses. The heatmap illustrates pairwise correlations between module eigengenes, highlighting relationships among structural remodelling, metabolic, bioenergetic, mechano-signalling, and regulatory transcriptional programs.

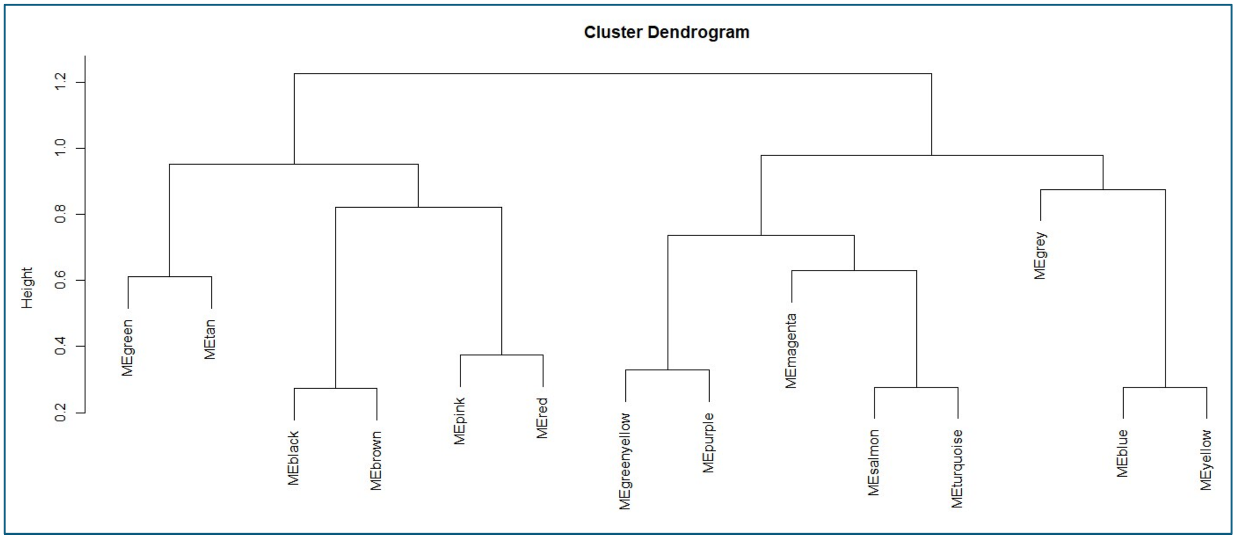


**Figure S4.** Cluster dendrogram of the 12 modules identified by WGCNA. We excluded the pink module because it was closely related to the Red module.

**Table S6.** Characteristics of the principal WGCNA modules

| Module | Number of genes | Correlation with LA weight (r) | P value |
|---|---|---|---|
| Turquoise | 3232 | 0.812 | $3.22 \times 10^{-9}$ |
| Blue | 2580 | -0.625 | $5.89 \times 10^{-5}$ |
| Red | 431 | -0.610 | $1.01 \times 10^{-4}$ |
| Black | 237 | 0.606 | $1.16 \times 10^{-4}$ |
| Pink | 184 | -0.511 | $1.70 \times 10^{-3}$ |
| Purple | 88 | 0.472 | $4.19 \times 10^{-3}$ |
| Salmon | 52 | 0.438 | $8.58 \times 10^{-3}$ |
| Green | 694 | 0.401 | $1.71 \times 10^{-2}$ |
| Tan | 75 | -0.182 | $2.96 \times 10^{-1}$ |
| GreenYellow | 84 | -0.157 | $3.67 \times 10^{-1}$ |
| Magenta | 139 | 0.115 | $5.11 \times 10^{-1}$ |
| Yellow | 1874 | -0.111 | $5.27 \times 10^{-1}$ |
| Brown | 2575 | 0.069 | $6.96 \times 10^{-1}$ |
| Grey | 565 | 0.001 | $9.94 \times 10^{-1}$ |

**Table S7**. A selection of genes modulated in a sex-specific manner in the Purple module.

| Gene | Male log2FC | Male FDR | Female log2FC | Female FDR | Biological function |
|---|---|---|---|---|---|
| Xirp1 | 1.46 | $1.68\times10^{-9}$ | -0.08 | 0.828 | Actin-binding protein involved in cardiomyocyte structural remodelling |
| Ccn1 | 1.26 | 0.0083 | 0.26 | 0.630 | Matricellular signalling protein regulating ECM remodelling and angiogenesis |
| Egr2 | 1.65 | 0.0456 | -0.76 | 0.083 | Immediate-early transcription factor involved in stress adaptation |
| Gadd45g | 1.28 | 0.0093 | -0.61 | 0.079 | Stress-response regulator controlling growth arrest and apoptosis |
| Arid5a | 1.09 | $4.79\times10^{-5}$ | 0.08 | 0.807 | RNA-binding transcriptional regulator involved in inflammatory signalling |
| Dnajb5 | 0.65 | 0.0100 | 0.11 | 0.758 | Molecular chaperone associated with protein quality control |
| Abra | 0.94 | 0.0061 | 0.10 | 0.674 | Actin-associated regulator of muscle growth and mechanotransduction |
| Nab2 | 0.74 | $2.0\times10^{-4}$ | 0.30 | 0.205 | Transcriptional coregulator downstream of early growth response pathways |
| Trib1 | 0.64 | 0.0056 | -0.03 | 0.955 | Pseudokinase involved in MAPK and metabolic signalling. |
| Ier5 | 0.75 | 0.0296 | -0.32 | 0.369 | Immediate-early response gene involved in cellular adaptation to stress |

**Table S8.** Persistent genes (number and % of total genes) within WGCNA modules.

| Module | Persistent genes | Biological identity |
|---|---|---|
| Blue | 133 (5%) | Mitochondrial metabolism |
| Turquoise | 93 (3%) | Structural remodeling |
| **GreenYellow** | **39 (46%)** | **Stress/adaptive response** |
| Purple | 5 (6%) | Regulatory signaling |

**Table S9.** Representative persistent genes within the green-yellow module.

| Gene | MHS log2FC | MR log2FC | Biological role |
|---|---|---|---|
| Dusp1 | -0.72 | -1.27 | MAPK signalling feedback regulator |
| Dusp5 | -0.53 | -0.65 | ERK/MAPK pathway regulator |
| Klf2 | -0.62 | -0.99 | Flow-responsive transcription factor |
| Klf4 | -0.36 | -0.99 | Stress-responsive transcription factor |
| Nr4a2 | -0.84 | -1.38 | Immediate early nuclear receptor |
| Apold1 | -0.80 | -1.78 | Endothelial stress-response gene |
| Tnfaip6 | -0.37 | -1.00 | Inflammation and ECM remodelling |
| Ret | -0.65 | -1.67 | Receptor tyrosine kinase signalling |
| Cebpb | -0.44 | -0.67 | Transcriptional regulator of stress responses |
| Errfi1 | -0.54 | -0.92 | Negative regulator of EGFR signalling |
| Sik1 | -0.88 | -1.11 | Stress-activated kinase signalling |
| Adamts1 | -0.49 | -0.61 | Extracellular matrix remodelling protease |
| S1pr1 | -0.35 | -0.31 | Sphingosine-1-phosphate signaling |
| Trib3 | +0.37 | +0.56 | Cellular stress and metabolic signalling |
| Id2 | +0.52 | +0.66 | Cell differentiation and fate determination |
